# An Activity-Based Nanosensor for Minimally-Invasive Measurement of Protease Activity in Traumatic Brain Injury

**DOI:** 10.1101/2022.12.26.521967

**Authors:** Julia A. Kudryashev, Marianne I. Madias, Rebecca M. Kandell, Queenie X. Lin, Ester J. Kwon

## Abstract

Current screening and diagnostic tools for traumatic brain injury (TBI) have limitations in sensitivity and prognostication. Aberrant protease activity is a central process that drives disease progression in TBI and is associated with worsened prognosis; thus direct measurements of protease activity could provide more diagnostic information. In this study, we engineered a nanosensor that releases a measurable signal into the blood and urine in response to activity from the TBI-associated protease calpain. Readouts from our nanosensor were designed to be compatible with ELISA and lateral flow assays, clinically-relevant assay modalities. In a mouse model of TBI, we demonstrated greater sensitivity of the nanosensor with the addition of targeting ligands to hyaluronic acid. In evaluation of mice with mild or severe injuries, our nanosensor identified mild TBI with a higher sensitivity than the clinical biomarker GFAP. This nanosensor technology allows for measurement of TBI-associated proteases without the need to directly access brain tissue, and has the potential to complement existing TBI diagnostic tools.

## INTRODUCTION

Traumatic brain injury (TBI) affects approximately 2.8 million people annually in the United States and leads to the hospitalization of over 200,000 patients per year.^1^ TBI is currently diagnosed in the clinic using a combination of behavioral testing (e.g., Glasgow Coma Scale) and medical imaging (e.g., MRI, CT).^2,3^ While these modalities are important tools for initial triage and for identification of patients who require surgical intervention, they are limited in their sensitivity for mild TBI and in their ability to predict long-term outcomes.^2,3^ Given these challenges and the heterogeneous nature of TBI, more tools are needed to address diagnostic and prognostic needs at different stages of disease.^2,4^

Recent research has identified endogenous breakdown products released into the blood and cerebrospinal fluid as biomarkers for TBI.^4,5^ These breakdown products are released from the damaged brain during secondary injury; thus, the measurement of these biomarkers can help identify processes that otherwise could not be captured by medical imaging. Moreover, biomarkers can be sampled through minimally-invasive collection of biofluids and can be quantified inexpensively and with higher throughput compared to medical imaging. The only FDA-cleared TBI diagnostic to date is based on measurements of the biomarkers GFAP and UCH-L1, brain-specific proteins that are shed from the brain into the blood. Measurements of GFAP/UCH-L1 are used to stratify TBI patients for CT imaging to identify operable lesions.^6^ More recently, these two biomarkers were found to be strong predictors for mortality and unfavorable outcomes in moderate to severe TBI, but their prognostic ability was limited in predicting recovery in mild TBI and in identifying patients with incomplete recovery.^7^ However, the existing TBI blood biomarkers that are FDA-cleared are associative; they correlate with disease but do not necessarily measure processes that directly contribute to disease.

As an alternative to the measurement of endogenous biomarkers, there is a potential to evaluate disease progression through the monitoring of protease activity. Proteases contribute to critical cellular processes such as signal transduction and protein processing in health,^8^ while in disease the ectopic expression or activation of proteases can drive pathological processes such as inflammation and extracellular matrix remodeling. In TBI, many of the endogenous biomarkers currently under investigation for diagnosis (e.g., breakdown products of GFAP, αII-spectrin and myelin basic protein) are protein fragments produced by ectopic protease activity including activity from the calcium-dependent protease calpain.^5^ Calpain is expressed within neurons, astrocytes, and endothelial cells in the brain, and it is ectopically activated within injured brain tissue after TBI as part of the secondary injury.^9–12^ Ectopic calpain activity is implicated in cellular death and worsened prognosis after TBI,^13–16^ and calpain is a target of investigational therapeutics for TBI.^17^ Thus, tools to measure calpain activity may useful in diagnosing injury progression, predicting patient prognosis, and informing precision treatments after TBI as a complement to the current standard of care.

Protease activity has been measured with activity-based nanosensors (ABNs) which produce quantifiable signals upon protease-mediated cleavage of a peptide substrate. ABNs are engineered from the ground up, so their components can be selectively tuned to optimize the sensitivity and specificity of measurements beyond that of endogenous biomarkers.^18,19^ This strategy has been successfully applied for the detection of pathological protease activity in cancer.^18–20^ In previous foundational work, we engineered an imaging-based ABN for calpain-1 activity as a tool to visualize spatial protease activation in TBI.^12^ This nanosensor accumulated within the injured brain tissue and produced fluorescent signal after systemic administration in a mouse model of TBI,^12^ and in a subsequent study we improved the sensitivity of this calpain-1 nanosensor by the addition of a targeting ligand that binds to the brain extracellular matrix component hyaluronic acid (HA).^21^ However, there are currently no technologies available to measure protease activity in TBI without directly probing brain tissue and thereby requiring an invasive tissue collection or sacrifice of the living organism.

Our goal in this work was to engineer an activity-based nanosensor for TBI (TBI-ABN) that generates signals that are accessible outside the brain to allow for minimally-invasive measurement of calpain-1 activity after TBI. In the work herein, we demonstrate that our TBI-ABN technology can produce signals in the blood and urine in response to injury in a mouse model of TBI. We evaluated valency-dependent impacts of adding HA targeting ligands to the TBI-ABN and identified an optimal targeting valency where sensitivity could be improved without a trade-off in specificity. The targeted TBI-ABN was then assessed for diagnostic efficacy in both female and male mice at two injury severities, where we observed sex-based differences in urine signal but not in blood signal. In both sexes of mice, the TBI-ABN was found to be more sensitive to mild injury compared to blood measurements of the clinical biomarker GFAP. Lastly, we show that readouts from our engineered TBI-ABN could be measured with clinically-relevant formats, ELISA and lateral flow assays. This technology enables measurement of molecular activity in TBI without the need to collect brain tissue, and thus it has potential both as a tool to study correlations between protease activity and long-term outcomes in TBI research and as a TBI diagnostic that complements existing modalities.

## RESULTS

### Engineering TBI-ABN for two detection modalities

In previous work we developed a FRET-based nanosensor for calpain-1 activity which could provide spatial information on calpain-1 activation in injured brain tissue.^12,21^ To enable minimally-invasive measurements of calpain-1 activity, we redesigned our nanosensor for measurement by both direct fluorescence and immunoassay quantification (**Figure 1**). Direct fluorescence measurement was included as a simple method to validate the engineering design of the nanosensor, while immunoassay quantification was included both for an increased dynamic range and sensitivity and as a format relevant to clinical diagnostics. This nanosensor contains a calpain substrate peptide from native substrate αII-spectrin^22^ flanked by a fluorescein (FAM) fluorophore and CPQ2 quencher FRET pair (Biotin-GGSGG-K(5FAM)-QEVYGAMP-K(CPQ2)-C-NH2). The FRET pair enabled fluorescence measurements and the biotin/FAM fragment allowed detection with immunoassays. To prolong circulation time in the blood and improve passive accumulation into the injured brain, the calpain substrate peptide was conjugated via a C-terminal cysteine to an 8-arm 40 kDa poly(ethylene glycol) (PEG) scaffold.^12^ The multi-arm PEG scaffold can carry multiple functional moieties, and PEG is a component of many FDA-approved materials;^23^ moreover, it has a hydrodynamic diameter of ∼10 nm^12^ which is both above the renal filtration limit of ∼5.5 nm^24^ and below the estimated pore size of the brain extracellular matrix.^25,26^ Nanoscale materials are able to passively accumulate across the injured vasculature into brain tissue in the first 24 hours after TBI due to an enhanced permeation and retention-like effect.^12,21,27–30^ The peptide was designed such that upon calpain-1 cleavage, the quencher remains attached to the PEG scaffold while the biotin/FAM-modified cleaved peptide (c-Peptide) is released (**Figure 2a**). The c-Peptide contains an N-terminal biotin connected to the FAM via a short peptide linker to create a heterobifunctional synthetic biomarker for downstream analysis.^31^ This biomarker can be quantified with sandwich immunoassays using α-FAM antibodies to capture the FAM end of the peptide and labeled streptavidin to detect the biotin end of the peptide.

**Figure 1.**
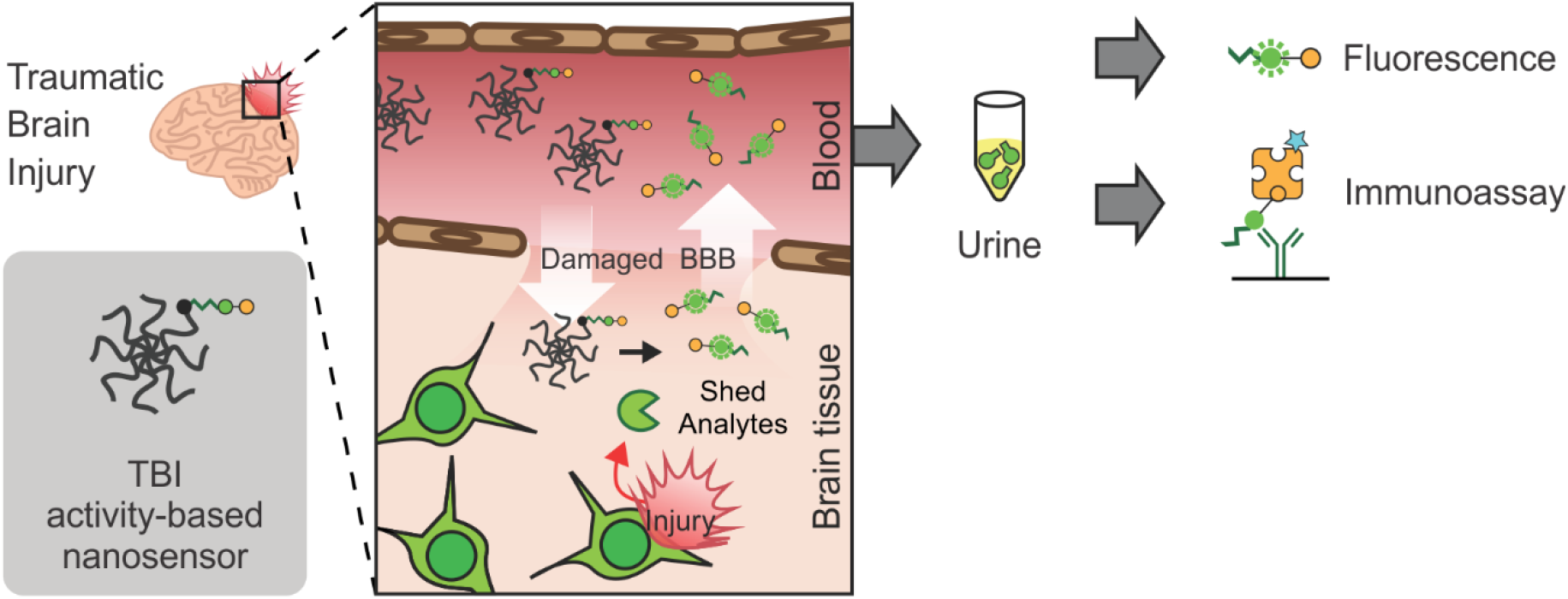
Overview of diagnostic TBI-ABN. After systemic administration, the ABN accumulates in injured brain tissue where it is cleaved by calpain-1. The cleaved peptide (c-Peptide) can then shed back into the blood and urine for minimally-invasive measurement of protease activity via fluorescence or immunoassay.

**Figure 2.**
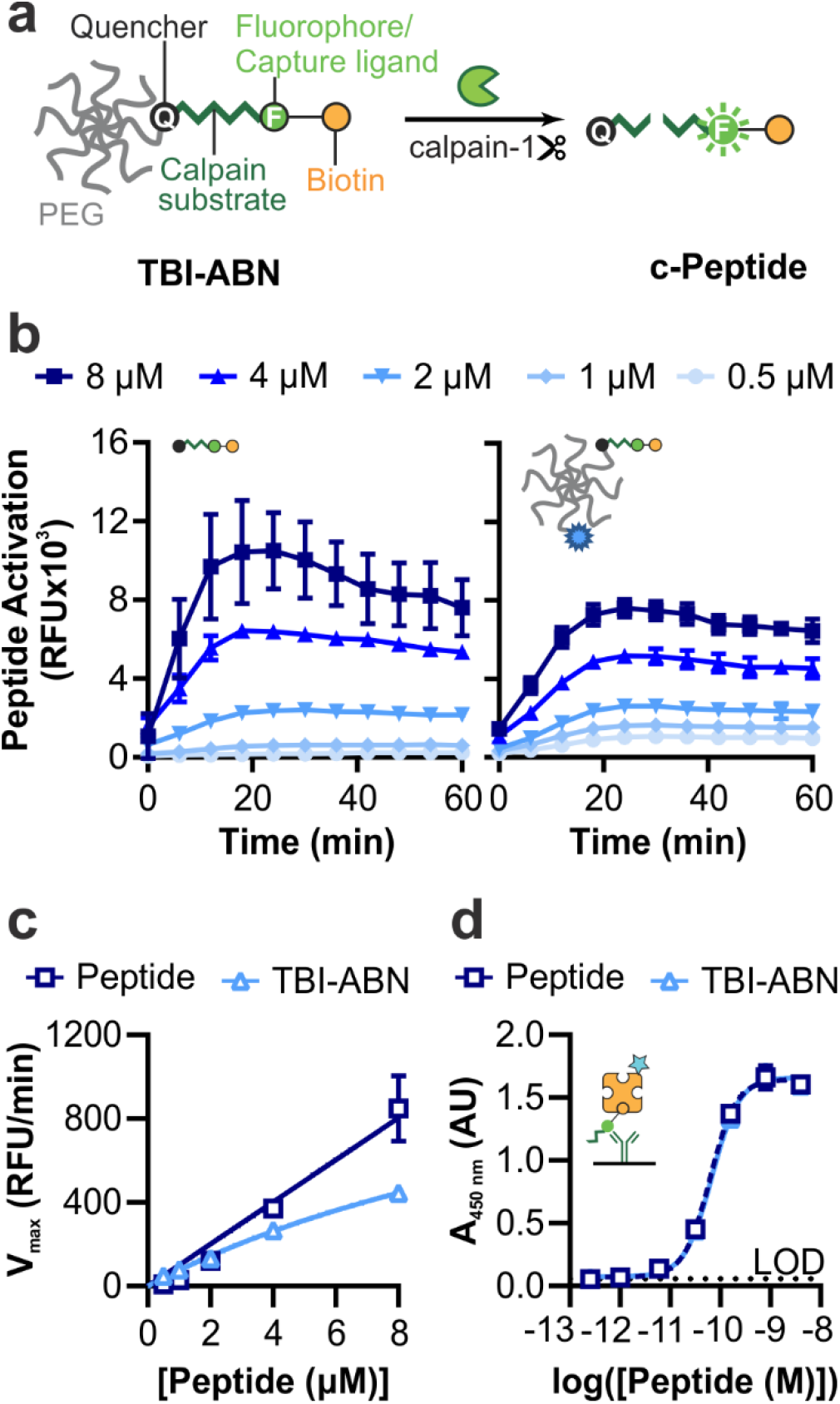
TBI-ABN contains a calpain-1 FRET substrate peptide that can be quantified via fluorescence and ELISA. **a**. Schematic of TBI-ABN, which releases a cleaved peptide (c-Peptide) biomarker after cleavage of the peptide substrate by calpain-1. **b**. In vitro cleavage kinetics of free peptide (left) and peptide conjugated to PEG to form TBI-ABN (right), incubated with human calpain-1 (n = 3 technical replicates, mean ± SD). **c**. Michaelis-Menten curves derived from the maximal velocities of cleavage (n = 3 technical replicates, mean ± SD). **d**. Dynamic range of free peptide and TBI-ABN on a sandwich ELISA, with α-FAM as the capture antibody and streptavidin-HRP as the detection molecule. (n = 3 technical replicates, mean ± SD, dashed line denotes the limit of detection (LOD)).

For initial kinetics and pharmacokinetics tests, TBI-ABN was synthesized by reacting 40 kDa 8-arm PEG-maleimide with 1 mole equivalent of calpain substrate peptide and 1 mole equivalent of VivoTag S-750 (VT750) fluorophore. We verified that the calpain substrate peptide undergoes FAM dequenching after cleavage by human calpain-1 in vitro, and that conjugation of the peptide to the PEG scaffold minimally impacted cleavage kinetics (**Figure 2b**). At 8 µM peptide, the maximal velocity of cleavage decreased from ∼850 RFU/min to ∼445 RFU/min with conjugation. However, at lower concentrations of peptide the difference in cleavage rates was less pronounced between free peptide and conjugate (**Figure 2c**). We investigated whether the decrease in maximal velocity was due to the presence of PEG, or due to the conjugation of peptide to PEG, and found that the conjugation of peptide to PEG led to the decrease in kinetics (**Figure S1**).

Previous studies have shown that the attachment of peptide substrates to PEG changes cleavage specificity and kinetics.^19^ The specificity of calpain substrate was assessed by incubating free peptide with two off-target proteases: α-thrombin, an abundant blood protease that is involved in the clotting cascade and can extravasate from the blood into the brain parenchyma in brain injury,^32–34^ and MMP9, a protease that has also been found to activate within injured brain tissue after TBI.^35–37^ The calpain substrate peptide did not activate after incubation with α-thrombin nor MMP9 at the concentrations tested (**Figure S2**). Lastly, to validate that the designed peptide could be sensitively measured in immunoassays, we developed a companion ELISA immunoassay. The ELISA uses α-FAM antibodies to capture the FAM and streptavidin-HRP to detect the biotin on the peptide. This allows for a sensitive measurement of calpain substrate peptide with a limit of detection of ∼1 pM (**Figure 2d**).

### TBI-ABN produces signal in the blood and urine in a TBI mouse model

The pharmacokinetics and activation of calpain substrate peptide and TBI-ABN were then evaluated in vivo in a mouse model of TBI using fluorescence as a direct readout of cleavage (**Figure 3a**). First, the blood circulation time and urine clearance were investigated to determine the time-dependent boundaries for sampling TBI-ABN activity. In uninjured mice, free peptide was intravenously administered followed by blood collection up to 2 hours post-injection or urine collection at 1 hour post-injection. Peptide was quantified via ELISA and compared to a standard to obtain the percent injected dose (%ID) for each sample. Free calpain substrate peptide was found to have a distribution half-life of 1.49 minutes, and a clearance half-life of 18.8 minutes (**Figure 3b**). At 1 hour post-injection, ∼5% of the peptide remained in blood circulation, while the greater majority of peptide was recovered from the urine (**Figure 3b-c**). This suggests that the peptide can efficiently clear into the urine after 1 hour of circulation, which was expected based on the small molecular weight of the peptide (∼2.6 kDa when uncleaved and ∼1.5 kDa when cleaved).

**Figure 3.**
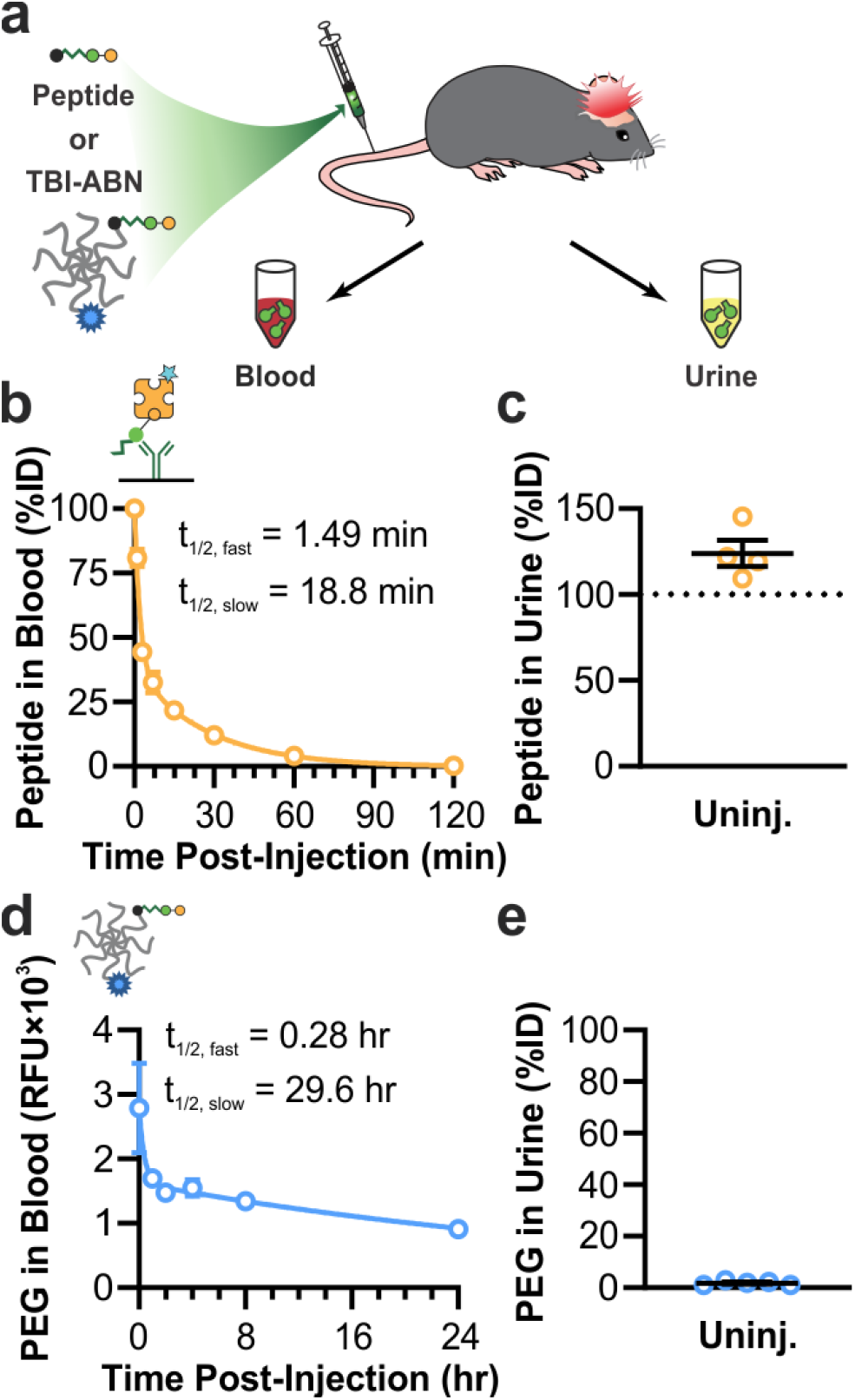
Pharmacokinetics of TBI-ABN in mice. **a**. Nanosensor was intravenously administered at 3 hours post-injury, and blood and urine samples were collected at multiple time-points after injection. **b**. Blood circulation half-life of free calpain substrate in the blood of uninjured mice, as measured by ELISA (n = 2 for 1 and 3 minutes; n = 4 for 7, 15, 30, 60, and 120 minutes; mean ± SE, two-phase decay). **c**. Estimated accumulation of free calpain substrate in the urine of uninjured mice at 1 hour post-injection, as measured by ELISA (n = 4, mean ± SE). **d**. Blood circulation half-life of TBI-ABN tagged with VT750 in uninjured mice, as measured by fluorescence of VT750 (n = 5, mean ± SE). **e**. Accumulation of TBI-ABN tagged with VT750 in urine of uninjured mice at 1 hour post-injection, as measured by fluorescence of VT750 (n = 5, mean ± SE).

To track the circulation of the larger TBI-ABN construct which is designed to remain in the blood within the sampling timeline, the PEG scaffold was conjugated with the near infrared fluorophore VT750. Uninjured mice were intravenously given TBI-ABNs followed by blood collection up to 24 hours post-injection and urine collection at 1 hour post-injection. Blood and urine samples were both measured for the fluorophore and compared to a standard to determine %ID. TBI-ABNs were found to have a much longer circulation time in the blood compared to free peptide, with a distribution half-life of 16.8 minutes and a clearance half-life of 29.6 hours (**Figure 3d**). At 1 hour post-injection, ∼1.8% of PEG was recovered from urine (**Figure 3e**). The observed clearance was comparable to previous observations on renal clearance of 40 kDa 8-arm PEG in mice,^18^ and is over an order of magnitude lower than the observed clearance of free peptide. The 40 kDa 8-arm PEG scaffold has a hydrodynamic diameter of ∼10.3 nm,^12^ which is above the renal filtration limit of ∼5.5 nm.^24^ Thus, the kidney acts as a natural size filter to allow c-Peptide to accumulate in the urine while excluding uncleaved, intact TBI-ABNs. This property allows for quantification of c-Peptide in the urine using both fluorescence and immunoassay without the need to separate uncleaved from cleaved TBI-ABN. Due to the inability to separate uncleaved and cleaved TBI-ABN in blood, TBI-ABN activation in the blood was only measured through fluorescence of activation. Due to the rapid rate of free peptide clearance, the accumulation of signal with injury is expected to take more time to be apparent in blood, which provides a snapshot of activity, compared to urine, which can integrate the accumulation of signal over time.^18^ The kinetics of nanosensor transport with the kinetics of nanosensor activation to optimize urine signal has been explored previously in a mathematical model of activity-based nanosensor activation in cancer.^18^

### Addition of hyaluronic acid targeting ligands improves sensitivity of TBI-ABN

Next, we sought to improve the sensitivity of blood and urine measurement by incorporating active targeting ligands to hyaluronic acid (HA). HA is highly abundant in the brain extracellular matrix and can be accessed via passive accumulation after disruption of the BBB in TBI.^21,27,38,39^ In our imaging-based nanosensor, we showed that the addition of a peptide ligand of HA (HApep)^40^ resulted in a 6.6-fold increase in signal over untargeted nanosensor.^21^ We hypothesized that this increased signal generation achieved with HA targeting in our imaging nanosensor would translate to increased signal in the blood and urine from our redesigned diagnostic TBI-ABN. The addition of targeting ligands was previously demonstrated to improve sensitivity of a cancer ABN.^19^

We investigated how the valency of targeting ligand impacted blood and urine signal generated from our diagnostic TBI-ABN. In order to create conjugates with matched levels of sensor peptide, 40 kDa 8-arm PEG-maleimide and calpain substrate peptide were first reacted in a single batch then split into 3 reactions with different inputs of HApep with an N-terminal cysteine (C-W-STMMSRSHKTRSHHV).^40^ The resulting conjugates for non-targeted, moderate targeted, and high targeted TBI-ABNs had molar ratios of 0, ∼2.5, and ∼3.9 HApep per calpain substrate, respectively, as measured by absorbance (**Figure S3**). The addition of HApep on the TBI-ABNs did not significantly affect cleavage kinetics of the calpain substrate peptide between conjugates (**Figure S4, S5**).

Conjugates were evaluated in a controlled cortical impact (CCI) model of TBI. CCI is a well-characterized model of TBI^41^ and has shown to recapitulate human-relevant features of TBI, such as locally increased calpain-1 activity within injured brain tissue.^11,12^ Uninjured mice and CCI-injured mice were each intravenously administered one of the three targeted TBI-ABNs at 3 hours post-injury, following previous observations in increased calpain-1 activity in injured brain tissue at that time-point.^12^ Blood collection was performed at 0, 1, 2, 4, and 8 hours post-injection and urine collection was performed at 1 hour post-injection (**Figure 4a**). In fluorescence readings of calpain substrate cleavage from blood samples, non- and moderate targeted TBI-ABNs showed an increased activation of signal in injured mice relative to the baseline activation seen in uninjured mice (**Figure 4b**). In contrast, the high targeted TBI-ABNs showed minimal differences in blood signal between injured and uninjured mice. For further analysis, we generated receiver operating characteristic (ROC) curves for each time point based on the ability to distinguish signal between uninjured and CCI-injured mice and calculated the area under the curve (AUC) as a measure of diagnostic performance (**Figure S6**). ROC statistics for this figure and subsequent figures can be found in **Tables S1 and S2**. Based on ROC AUC values, the ability of each TBI-ABN to classify injury through blood samples increased over time from 1 hour through 4 hours before plateauing (**Figure 4c**). The moderate targeted TBI-ABN outperformed the non-targeted TBI-ABN at classifying injury through blood samples from 1 hour post-injection, while the high targeted TBI-ABN could only classify injury from 4 hours post-injection.

**Figure 4.**
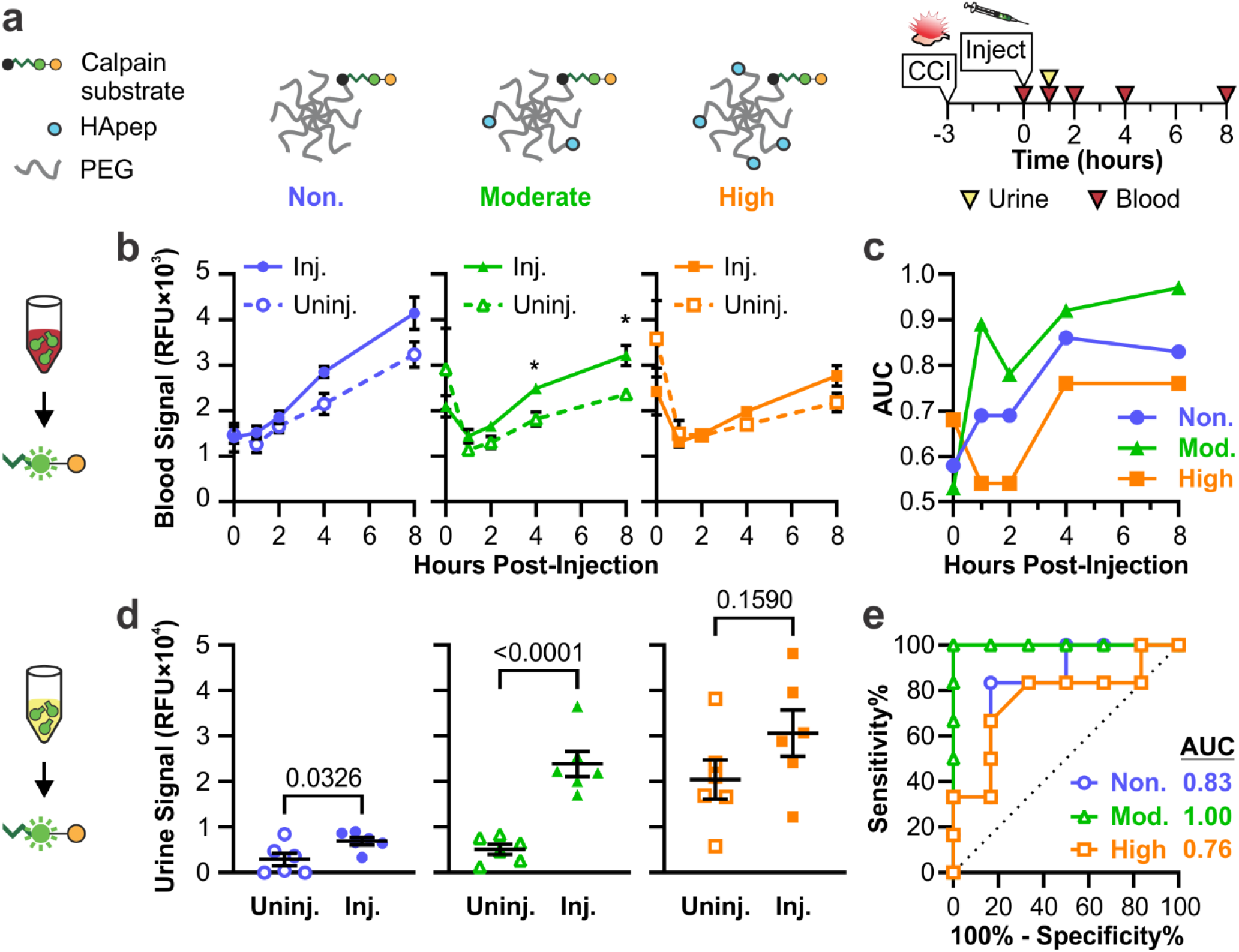
Targeting to hyaluronic acid improves the sensitivity of TBI-ABN in both blood and urine. **a**. Schematic of HA-targeted TBI-ABNs with non-, moderate, and high targeted ABNs. **b**. Activation of sensor in the blood of injured and uninjured mice at multiple time-points post-injection as measured by fluorescence (n = 6 per group, mean ± SE, Two-way RM ANOVA with Geisser-Greenhouse correction and Sidak’s multiple comparisons post-hoc test for each time point, *p<0.05). **c**. AUCs for blood ROCs taken at each time point (n = 6 per group). **d**. Accumulation of activated sensor in the urine of mice 1 hour post-injection as measured by fluorescence (n = 6 per group, mean ± SE, Unpaired t-test). **e**. ROC curves and calculated AUCs from fluorescence measurements of urine (n = 6 per group). See **Table S1** for additional ROC statistics.

Consistent with measured blood activation, the moderate targeted TBI-ABN showed the greatest increase in urine signal between injured and uninjured conditions compared to the non- and high targeted TBI-ABNs (**Figure 4d**). Interestingly, while the absolute fluorescent signal in urine increased with ligand valency in injured mice, high targeted TBI-ABN yielded substantially elevated signal accumulation in the urine of uninjured mice. The moderate targeted TBI-ABNs yielded the highest signal-to-noise ratio of 4.7, significantly outperforming the signal-to-noise ratios of 2.4 and 1.5 for non- and high targeted TBI-ABNs, respectively. These observations are consistent to previous observations with targeting with the imaging nanosensor,^21^ where the sensitivity of nanosensor signal within injured brain tissue was the highest for the high targeted nanosensor, at the cost of high accumulation and activation in off-target organs. In that study, the moderate targeted nanosensor yielded the highest specificity with minimal accumulation or activation within off-target organs. The specificity advantage of moderate levels of targeting were recapitulated in the diagnostic TBI-ABN; moderate targeted TBI-ABN was able to discriminate injury with an ROC AUC of 1.00 (95% confidence interval (CI) 1.00-1.00), compared to non-targeted and high targeted TBI-ABNs with ROC AUCs of 0.83 (0.59-1.00) and 0.76 (0.47-1.00), respectively (**Figure 4e**). The 4-hour post-injection (7-hour post-injury) time point was selected for blood sampling in subsequent experiments based on the magnitude of difference observed in the time course of injured signal relative to uninjured controls (**Figure 4b**), and to be compatible with clinically-relevant time-frames for blood collection in TBI.^6,42,43^

### Sex- and severity-dependent performance of targeted TBI-ABN

After establishing the optimal valency of HA targeting and identifying the time points for blood and urine collection, we further evaluated the diagnostic efficacy of the moderate targeted TBI-ABN in a larger cohort of female and male mice and two injury severities. Sex-based differences in cellular responses and outcome after TBI have been observed in both human patients and animal models,^44,45^ and these differences could extend to calpain activity. For example, the spatial distribution and time course of calpain activation in the brain was found to differ after diffuse head injury in female and male mice.^46^ Thus, we investigated whether the TBI-ABN could detect sex-based differences in calpain activation. We also investigated whether the TBI-ABN could discriminate between different severities of injury, as the serum levels of calpain-specific breakdown products have previously been observed to increase with severity of injury in human patients.^15^ Two severities of injury were investigated based on CCI parameters commonly used in TBI research.^47^ For mild TBI, impact depth was 0.5 mm and impact speed was 1 m/s; for severe TBI, impact depth was 2 mm and impact speed was 5 m/s. TBI-ABN was administered at 3 hours post-injury, urine was collected at 1 hour post-injection (4 hours post-injury), and blood was collected at 4 hours post-injection (7 hours post-injury) (**Figure 5a**).

**Figure 5.**
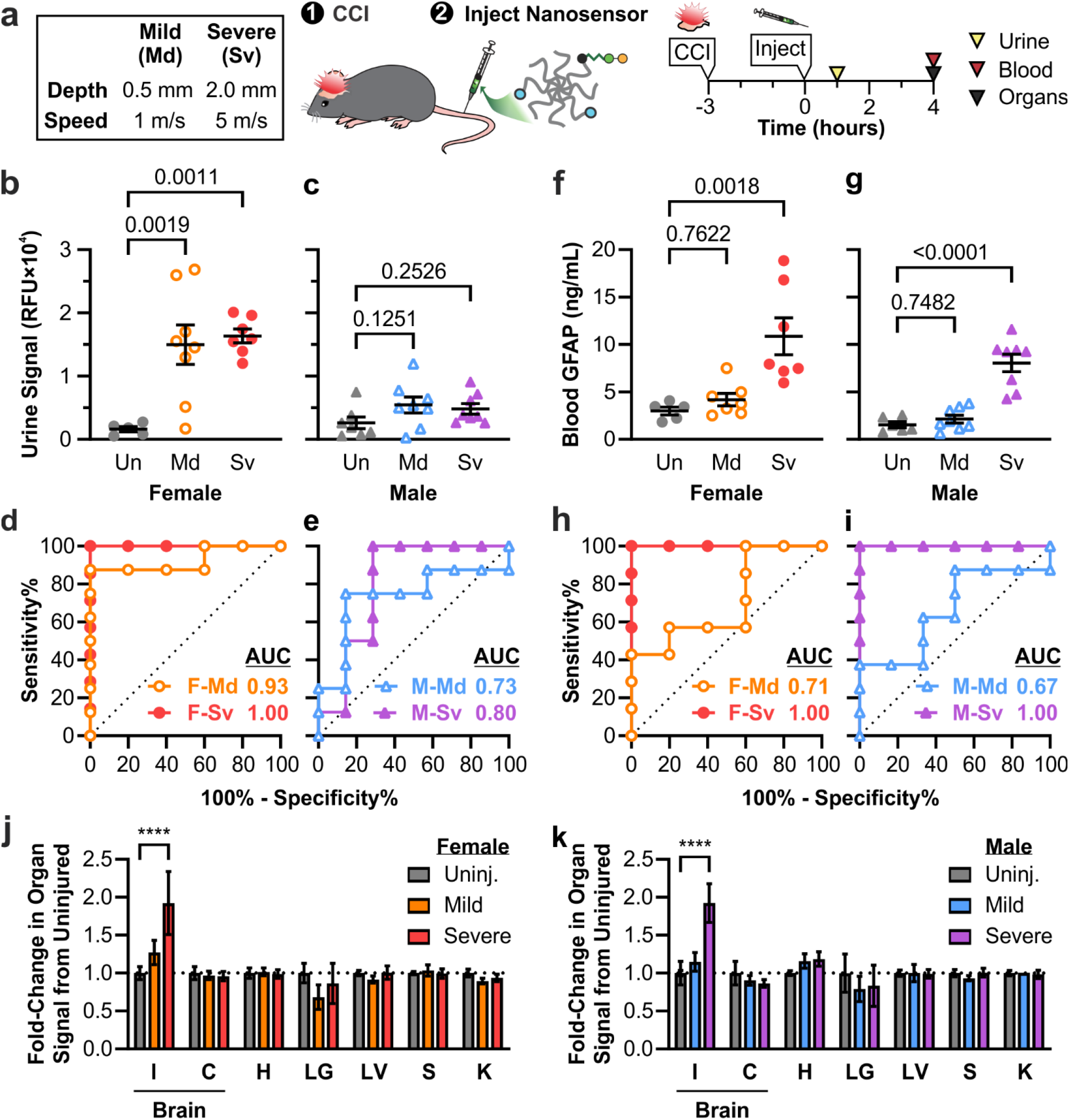
Moderate targeted TBI-ABNs show different patterns of signal with injury in female vs. male mice. **a**. Injury conditions in mice and experimental timeline. **b, c**. TBI-ABN activation in urine at 1 hour post-injection as measured by fluorescence and **d, e**. corresponding ROC curves in female and male mice, respectively (Un = uninjured; Md = mild CCI; Sv = severe CCI; F = female; M = male; n = 5 for F-Un; n = 7 for F-Sv and M-Un; n = 8 for F-Md, M-Md, and M-Sv; mean ± SE, ordinary one-way ANOVA with Dunnett’s multiple comparisons post-hoc test against uninjured control). **f, g**. GFAP levels in blood at 4 hours post-injection as measured by ELISA and **h, i**. corresponding ROC curves in female and male mice, respectively (n = 5 for F-Un; n = 6 for M-Un; n = 7 for F-Md and F-Sv; n = 8 for M-Md and M-Sv; mean ± SE, ordinary one-way ANOVA with Dunnett’s multiple comparisons post-hoc test against uninjured control). **j, k**. Relative fold-change of TBI-ABN activation in the brain (I = injured cortical tissue, C = contralateral cortical tissue, with I+C pooled for uninjured control) and in off-target organs (H = heart; LG = lungs; LV = liver; S = spleen; K = kidney) in female and male mice, respectively, with mild or severe CCI compared to uninjured controls (n = 5 for F-Un; n = 7 for F-Sv and M-Un; n = 8 for F-Md, M-Md, and M-Sv; mean ± SE, two-way ANOVA with Dunnett’s multiple comparisons test compared to uninjured within each organ, ****p<0.0001). See **Table S2** for additional ROC statistics.

In urine samples collected at 1 hour post-injection, female mice produced strong fluorescent signals with both mild and severe injury, with a ∼9-10-fold increase in signal compared to uninjured female mice (**Figure 5b**). Curiously, male mice produced less fluorescent signal in urine with injury, showing a ∼2-fold increase in signal relative to uninjured male mice (**Figure 5c**). No significant differences in signal magnitude were observed between mild and severe injury conditions within either sex, although signals for mild injuries were more variable. The TBI-ABN was able to discriminate injury in female mice with strong diagnostic performances for both severe injury (AUC 1.00, CI 1.00-1.00) and mild injury (AUC 0.93, CI 0.77-1.00) (**Figure 5d, Table S2**). In male mice, the diagnostic performance was weaker, with ROC AUCs of 0.80 (0.55-1.00) for severe injury and 0.73 (0.46-1.00) for mild injury (**Figure 5e**). In blood samples taken 4 hours post-injection, the measured sensor signal (**Figure S7a-b**) and diagnostic performance (**Figure S7c-d**) was similar in female and male mice across injury severities, with ROC curve AUCs ranging from 0.77-0.89 depending on sex and injury severity.

To benchmark the diagnostic performance of our TBI-ABN, blood plasma samples at 7 hours post-injury were also quantified for GFAP concentration via ELISA. GFAP is one of the leading biomarkers that has been cleared by the FDA to determine the need for a CT scan after TBI, and its blood levels are known to increase within the first 12-24 hours after injury;^6,42,43^ thus the 7-hour blood time-point was selected to be within these clinically-relevant time frames. Blood levels of GFAP were found to be significantly elevated in both female and male mice with severe injury (**Figure 5f-g**), and could discriminate severe injury with an ROC of 1.00 (1.00-1.00) in both sexes (**Figure 5h-i**). Interestingly, blood levels of GFAP were only slightly increased with mild injury and had a weaker discriminatory ability than in severe injury, with ROC AUCs of 0.71 (0.41-1.00) and 0.67 (0.37-0.96) for female and male mice, respectively. Among TBI-ABN measurements for severe injury, urine fluorescence measurements from female mice at 4 hours post-injury matched the strong diagnostic efficacy of blood GFAP levels at 7 hours post-injury (**Figure 5d, h**). Moreover, the TBI-ABN outperforms GFAP in mild injury conditions, especially for urine measurements in female mice (**Figure 5d, h**).

To determine whether the observed sex-based differences in urine signal were due to differences in overall sensor activation in the injured brain, cortical brain tissue and off-target organs were collected for homogenization from each mouse at 4-5 hours post-injection. Sensor activation was measured from organ homogenates by fluorescence, as we have reported previously.^21^ A clear severity-dependent increase in fluorescence was observed only in the ipsilateral cortical brain tissue of injured mice and not in off-target organs, indicating that elevated urine signal was driven by the presence of brain injury (**Figure 5j-k, S8**). Tissue-level nanosensor activation was comparable between female and male mice within injury conditions. This suggests that tissue-driven sensor activation is independent of sex, and that the sex-based differences measured in urine signals could be due to differences in transport of c-Peptide rather than differences in signal generation.

### Measurement of c-Peptide in urine samples via immunoassays

To demonstrate that our TBI-ABN platform was compatible with clinically-relevant immunoassays, urine samples collected in **Figure 5a** were measured with an ELISA. The c-Peptide concentration quantified by ELISA correlated closely with fluorescence measurements with a Pearson correlation of r = 0.9313, confirming that an immunoassay accurately reflected peptide cleavage as measured by the direct fluorescent measurement (**Figure S9**). The ELISA measurements resulted in signal-to-noise ratios of ∼4.2-4.4 for injured female mice and ∼1.8-1.9 for injured male mice, respectively (**Figure 6a-b**). The ROC curves based on ELISA measurements likewise were comparable to the ROC curves based on fluorescence, with ROC AUCs of 0.98 (0.90-1.00) and 1.00 (1.00-1.00) for female mice with mild and severe injury, and ROC AUCs of 0.73 (0.45-1.00) and 0.87 (0.66-1.00) for male mice with mild and severe injury, respectively (**Figure 6c-d**).

**Figure 6.**
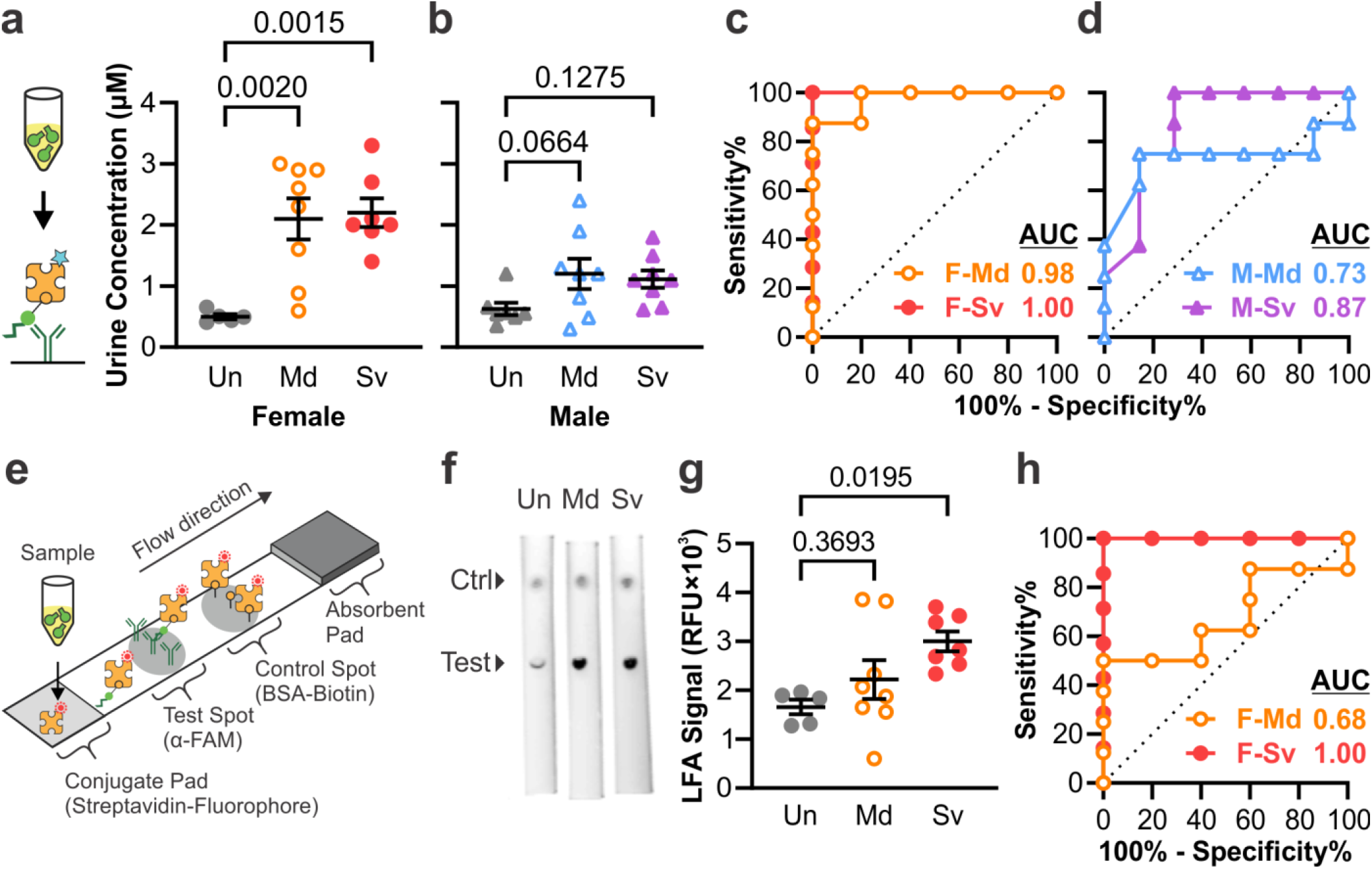
TBI-ABNs can be quantified from urine samples via immunoasssay. **a, b**. ELISA measurement of c-Peptide accumulation in urine at 1 hour post-injection and **c, d**. corresponding ROC curves in female and male mice, respectively. (Un = uninjured; Md = mild CCI; Sv = severe CCI; F = female; M = male; n = 5 for F-Un; n = 7 for F-Sv and M-Un; n = 8 for F-Md, M-Md, and M-Sv; mean ± SE, ordinary one-way ANOVA with Dunnett’s multiple comparisons post-hoc test against uninjured control). **e**. Schematic of LFA developed to measure c-Peptide in samples. **f**. Representative fluorescence scans of LFAs for urine samples from female mice (ctrl = control). **g**. Accumulation of c-Peptide in urine at 1 hour post-injection as measured by integrated test spot signal from each LFA and **h**. corresponding ROC curves in female mice (n = 5 for F-Un; n = 7 for F-Sv; n = 8 for F-Md; mean ± SE, ordinary one-way ANOVA with Dunnett’s multiple comparisons post-hoc test against uninjured control). See **Table S2** for additional ROC statistics.

A lateral flow assay (LFA) was also developed as another format of immunoassay that can detect biomarkers more rapidly than ELISA (3 hours vs. 20 min), although with a trade-off in assay sensitivity. This assay measures c-Peptide in a sandwich format using the same capture and detection ligands as in the ELISA (**Figure 6e**). α-FAM antibody was spotted onto a nitrocellulose membrane to capture the FAM on the c-Peptide, and fluorescently-labeled streptavidin was dried onto a conjugate pad for detection of the biotin on the c-Peptide. BSA-biotin was spotted distal to the test spot as a positive control. LFAs were tested in a dipstick format by dipping the conjugate pad end of the LFA into diluted urine samples. In a dilution standard of free peptide spiked into diluted mouse urine, the LFA could detect peptide over a range of 3.9-250 nM (**Figure S10**). Urine samples from female and male mice with mild and severe injury from **Figure 5a** were then applied to the LFA to determine whether the assay could stratify by injury (**Figure 6f and S11**). Urine samples from female mice with severe injury had sufficiently increased signal to allow stratification from uninjured mice with an ROC AUC of 1.00 (1.00-1.00) (**Figure 6g-h**). As expected, the LFA showed less difference in urine samples for female mice with mild injury or in male mice with either injury severity, and thus had weaker diagnostic ability with ROC AUCs of 0.55-0.68 (**Figure 6g-h and S12**). The LFA format allows for rapid measurement of urine signal with the trade-off of lowered sensitivity.

## DISCUSSION

In summary, we have engineered a TBI-ABN that can identify brain injury via minimally-invasive measurements from the urine. We optimized the TBI-ABN for sensitivity and specificity through the addition of hyaluronic acid targeting ligands. We evaluated the diagnostic efficacy of our TBI-ABN in female and male mice across two severities of TBI. The TBI-ABNs were able to identify mild and severe injuries through urine measurements from female mice. Interestingly, urinary measurement of our TBI-ABN did not perform as well in male mice despite comparable tissue-level activation in both male and female mice. Moreover, the TBI-ABN appeared to be more sensitive to mild injury compared to blood GFAP levels across both sexes.

The observed difference in diagnostic performance between the nanosensors with various targeting valencies (**Figure 4**) highlights the importance of tuning targeting valency in nanomaterial interactions. The overall binding strength of nanomaterials to their targets can be enhanced by increasing targeting valency.^48^ However, too high of an avidity can disrupt transport of materials to their desired target,^49^ and changes in pharmacokinetic properties such as charge can affect cellular uptake in off-target organs such as the liver, spleen, or kidneys.^30,50^ This was observed with HA-targeting of a calpain-1 nanosensor in our previous study, where increasing valencies of targeting led to increased nanosensor activation within both the target brain tissue and in off-target organs.^21^ Thus, the relative contribution of off-target activation is likely increased with increasing targeting valency, leading to loss of specificity.

In the context of TBI, it has previously been shown that there are sex-dependent differences in the extravasation of nanoparticles to the injured brain 24 hours after TBI, but not at 3 hours after TBI.^51^ The lack of sex-based differences in nanosensor activation within injured brain tissue (**Figures 5j-k and S8**) and the shorter time-frame of this study both suggest that the observed differences in urine signal (**Figure 5b-c**) were not due to sex-dependent changes in nanosensor extravasation. The difference in transport could instead be due to intrinsic differences in renal function between sexes. Sex-based differences in the renal transport of different compounds, and in the expression of peptide transporters within the kidneys have been identified within multiple animal models and within humans.^52–55^ Initial optimization of the TBI-ABN (**Figures 2-3**) was done in female mice, and it’s likely different optimization parameters would be found in male mice. These observations bring up interesting questions on sex-dependent changes in physiology after TBI, which are out of scope of the current study.

While our work established the ability to measure protease activity in TBI from liquid biopsies, further work can be done to optimize diagnostic performance of the TBI-ABN. For example, the speed of signal detection in the blood could be increased by using substrate sequences that have higher calpain cleavage rates.^56^ The rate of c-Peptide urine clearance could also be increased by incorporating ligands optimized for renal clearance, such as glutamate-fibrinopeptide B or cyclodextrin.^57,58^ Finally, it is known that calpain activation occurs across other diseases such as Alzheimer’s and ischemia-reperfusion injury;^59–61^ the specificity of TBI detection can be enhanced by multiplexing the technology for additional proteases that are activated in TBI, such as MMP9 and cathepsin B.^35–37,62,63^ All of these parameters should be considered in the context of different demographic groups to ensure that the sensor can work in as many patients as possible.

Beyond initial diagnosis, protease activity measurements have the potential to aid in the evaluation of therapeutics for TBI. To date there are still no neuroprotective therapies for TBI, and one reason for failed clinical trials has been the lack of adequate quantitative biomarkers to assess therapeutic response.^64^ Recent work in the Operation Brain Trauma Therapy screening consortium has included biomarkers such as blood levels of GFAP in the evaluation of therapeutics, although there are still challenges in linking changes in these associative biomarker levels to the direct engagement of therapeutic targets.^65^ Proteases including calpain remain as potential targets for therapeutics,^17^ and thus tools to directly measure protease activity could be used to identify patients that have elevated calpain activity and monitor therapeutic response in a precision medicine strategy. Precision medicine has already been applied to drug discovery and disease monitoring in diseases including cancer and infection,^20,66,67^ and has the potential to supplement the diagnostic toolbox for TBI.

## METHODS

### Synthesis of TBI-ABN Poly(ethylene glycol) (PEG) Conjugates

Calpain substrate FRET peptide (Biotin-GGSGG-K(5FAM)-QEVYGAMP-K(CPQ2)-C-NH2) was synthesized by CPC Scientific Inc. (Sunnyvale, CA). HApep (C-W-STMMSRSHKTRSHHV) was synthesized by Lifetein (Somerset, NJ). The 40kDa 8-arm PEG-maleimide (tripentaerythritol) was purchased from Jenkem Technology (Beijing, China). For the initial kinetics and pharmacokinetics studies: PEG-maleimide was reacted with 1 mol equivalence of calpain substrate peptide and 1 mol equivalence of L-cysteine-functionalized VivoTag S-750 (PerkinElmer, Boston, MA), followed by quenching with an excess of L-cysteine. For the targeting comparison and diagnostic evaluation: PEG-maleimide was batch reacted with 1 mol equivalence of calpain substrate peptide to ensure matched substrate peptide modification for each targeting group. The reaction was then split and reacted with 0 mol (non-targeted), 4 mol (moderate targeted), or 7 mol (high targeted) equivalencies of HApep followed by quenching with an excess of L-cysteine. All conjugates were dialyzed with water, and final concentrations were determined by absorbance of VivoTag 750, FAM, or tryptophan using a Spark multimode microplate reader (Tecan Trading AG, Switzerland).

### In Vitro Reaction Kinetics Assay

Protease reaction kinetics assays were run as previously described.^12^ Briefly, calpain substrate peptide or conjugates were incubated with 26.6 nM human calpain-1 (Sigma-Aldrich, C6108) in 50 mM N-(2-hydroxyethyl)piperazine-N′-ethanesulfonic acid (HEPES), 50 mM NaCl, 2 mM EDTA, 5 mM CaCl_2_, and 5 mM β-mercaptoethanol. For specificity reactions, calpain substrate peptide was incubated with 15 nM human recombinant MMP-9 (Enzo Life Sciences, BML-SE360-0010) or 13.5 nM human α-thrombin (Haematologic Technologies, HCT-0020) in TCNB buffer. Fluorescence readings were taken every 90 s at 37 °C for 1 h using a Spark multimode microplate reader. Reaction curves were normalized to controls, and their initial velocities were fitted to a Michaelis−Menten curve in GraphPad Prism® (9.4.1).

### Controlled Cortical Impact (CCI) TBI Mouse Model

All mouse procedures were approved by the University of California San Diego’s Institutional Animal Care and Use Committee (IACUC). Female C57BL/6J mice (Jackson Labs) at 9-13 weeks old were used for all in vivo studies. Male C57BL/6J mice (Jackson Labs) at 9-13 weeks old were used in the last diagnostic evaluation. Following anesthetization with 2.5% isoflurane, buprenorphine analgesia was administered. A 5 mm craniotomy was performed over the right hemisphere between bregma and lambda, and controlled cortical impact was performed using the ImpactOne (Leica Biosystems) with a 2 mm diameter stainless steel probe and a dwell time of 300 ms. For the targeting comparison study, severe CCIs were performed with a velocity of 5 m/s and impact depth of 2 mm. In the diagnostic evaluation, an additional mild CCI condition was included with a velocity of 1 m/s and impact depth of 0.5 mm. The center of the injury impact was centered around -2.0 mm (± 0.5 mm) lateral from the midline and -2.0 mm (± 0.5 mm) caudal from bregma.

### Peptide and PEG Half-Life and Urine Accumulation Studies

To assess the pharmacokinetics of calpain substrate peptide, uninjured mice were administered peptide in 1xPBS at a dose of 50 nmol calpain substrate / kg through the tail vein. In one set of mice, blood was collected at 1, 3, 7, 15, 30, 60, and 120 minutes post-injection for quantification via ELISA (n = 2 for 1 and 3 minutes; n = 4 for 7, 15, 30, 60, and 120 minutes). In a separate group of mice, urine was collected at 1 hour post-injection for quantification via ELISA (n = 4). For the percent injected dose (%ID) of peptide, blood volume was estimated to be of 8% / kg. Blood half-life was estimated with two-phase decay after assuming 100 %ID at 0 minutes. Urine volume was estimated based on urine weight with a specific gravity of 1.0058. Percent accumulation of peptide into urine was estimated by multiplying the quantified concentration of peptide in urine by the volume to obtain the moles recovered, then dividing that value by the injected dose per mouse. To assess PEG biodistribution, uninjured mice were administered fluorescently-tagged TBI-ABN in PBS at 50 nmol calpain substrate / kg through the tail vein (n = 5). Urine was collected at 1 hour post-injection to assess PEG accumulation via fluorescence of VT750. Blood was collected at 0, 1, 2, 4, 8, and 24 hours post-injection and measured for PEG via fluorescence of VT750 using an Odyssey scanner (Li-Cor Biosciences). Percent accumulation of PEG into urine was calculated similarly to the calculation for percent accumulation of peptide, except that the concentration of PEG was based on a fluorescence standard of TBI-ABN.

### TBI-ABN In Vivo Studies

For comparison of targeted TBI-ABNs, non-, moderate, or high targeted TBI-ABN in 1xPBS was administered through the tail vein at 50 nmol calpain substrate / kg and at 3 hours post-injury. Uninjured mice and mice with severe CCI were compared for each targeting group (n = 6). Urine was collected at 1 hour post-injection and blood was collected at 0, 1, 2, 4, and 8 hours post-injection for quantification of TBI-ABN activation via FAM fluorescence. In the final diagnostic evaluation, female and male mice were compared with uninjured mice, mice with mild CCI, and mice with severe CCI for each sex (n = 5 for female uninjured; n = 7 for female severe and male uninjured; n = 8 for female mild, male mild, and male severe). Moderate targeted TBI-ABNs were administered at 50 nmol calpain substrate / kg through the tail vein at 3 hours post-injury. Urine was collected at 1 hour post-injection and blood was collected at 4 hours post-injection for quantification of TBI-ABN activation via FAM fluorescence. At 4-5 hours post-injection, mice were transcardially perfused with ice cold PBS and the organs were collected for homogenization to assess biodistribution of activated TBI-ABN.

### Blood Collection and TBI-ABN Measurement

At each time point, 10 µL of blood was collected from the tail vein into EDTA-coated tubes, then diluted 1:5 in PBS with a final concentration of 2 µM EDTA. Blood samples were centrifuged at 2,000 x g at 4 °C for 15 minutes to pellet cells and platelets, then the plasma supernatant was collected into clean tubes. Samples were immediately measured for fluorescence of FAM before storing at -80 °C. For blood pharmacokinetics of free peptide, blood samples were quantified via c-Peptide ELISA at a final dilution of 1:12,500 in PBS. For GFAP measurement via ELISA, blood samples were measured at a final dilution of 1:16.6 in Sample Diluent NS (Abcam).

### Urine Collection and TBI-ABN Measurement

After TBI-ABN injection, mice were given 200 µL subcutaneous USP saline to induce urine production, then placed in a urine collection box for 1 hour. After 1 hour, the bladders were gently pressed to expel all available urine and the mice were returned to their housing. Collected urine was weighed to estimate volume, then spun down at 10,000 x g at 4 °C for 2 minutes to pellet solid particles. Clean supernatant was transferred to a clean tube, diluted 1:25 in PBS for fluorescence measurement, then stored at - 80 °C. For c-Peptide ELISA measurement, urine samples were measured at a final dilution of 1:50,000 in PBS. For LFA measurement, urine samples were measured at a final dilution of 1:40 in PBS, 0.05% Tween-20, and 1% BSA.

### Quantification of Peptide from Urine and Blood via ELISA

To quantify c-Peptide, 96-well plates were coated overnight with polyclonal α-FAM antibody (Invitrogen, A-11095) at 0.5 µg/mL in PBS. Coated plates were then blocked with blocking buffer (PBS, 0.05% Tween-20, 1% BSA) for 2 hours, incubated with sample in blocking buffer for 1 hour, and incubated with 1:200 streptavidin-HRP (R&D Systems, DY998) for 1 hour. In between each incubation, the plate was washed with PBS with 0.05% Tween-20. TMB substrate (Thermo Scientific, 34021) was then added to develop signal for 5-10 minutes, followed by quenching with 1 N HCl. The final plate was quantified by absorbance at 450 nm. For the half-life study, blood and urine samples were quantified against a dilution ladder of injected material in PBS. For the targeting study, urine samples were quantified against a dilution ladder of free peptide in PBS. To quantify GFAP levels in blood, a Mouse GFAP SimpleStep ELISA kit was used (Abcam, ab233621). Manufacturer protocols were followed with the exception that Sample Diluent NS was substituted in place of Cell Extraction Buffer.

### Lateral Flow Assay

For the test and control spots, 0.3 µL of 1 mg/mL α-FAM antibody (Invitrogen, A-11095) or 0.3 µL of 50 ug/mL BSA-biotin, respectively, were spotted onto a nitrocellulose membrane (Cytiva, FF170HP) and dried for 30 minutes. A conjugate pad (Ahlstrom, A6614) was soaked in a solution of 30 µg/mL IRDye 800CW Streptavidin (Bio-Rad), 2.5% BSA, 10% sucrose, and 1% Tween-20 in PBS, then dried at 37 °C for 2 hours. The conjugate pad, membrane, and an absorbent pad (Millipore CFSP223000) were assembled on a backing card (DCN MIBA-020) then cut into 3-mm-wide strips and stored at room temperature. Dilution standards of free peptide were prepared in sample diluent buffer (PBS, 0.05% Tween-20, 1% BSA) spiked with a 1:40 dilution of mouse urine. Urine samples were diluted 1:40 in sample diluent buffer, then applied in 40 µL volumes to each assay in a dipstick format and run for 20-30 minutes. LFAs were imaged on an Odyssey scanner, and test spot signals were quantified in ImageJ (1.53t) via densitometry.

### Quantitative Biodistribution of Sensor Activation in Homogenized Tissue

Homogenization was performed as previously described.^21^ The collected organ tissue was flash frozen at -80 °C. The heart, lungs, spleen, and kidney were minced, and homogenization buffer (6% w/v sodium dodecyl sulfate (SDS), 150 mM Tris-HCl pH 7.4, 100 mM dithiothreitol (DTT), and 2 mM ethylenediaminetetraacetic acid (EDTA)) was added to all organs to a final concentration of 250 mg tissue / mL buffer. Tissue was further processed with a Tissue-Tearor with a 4.5 mm probe (Fisher) until lysate was fully homogenized. Samples were heated at 90 °C for 10 minutes with agitation at 800 RPM, vortexed to mix, and homogenate measured for FAM fluorescence to quantify relative activation of TBI-ABN.

### Software and Statistics

GraphPad Prism® (9.4.1) was used to perform statistics. All t-tests and post-hoc tests were conducted with an alpha of 0.05 to identify statistical significance between samples. P values were adjusted for multiplicity for multiple comparisons. In results for the GFAP ELISA, outliers were identified and removed using the ROUT method with a maximum false discovery rate of 1%. For t-values, F-values, and degrees of freedom, see **Table S3**.

## ACKNOWLEDGEMENTS

This work was supported by a National Science Foundation (NSF) CAREER Award (2046926) and the National Institutes of Health (NIH) Director’s New Innovator Award (DP2 NS111507). J.A.K. acknowledges support from the National Science Foundation (NSF) Graduate Research Fellowship Program under Grant No. DGE-1650112. Any opinions, findings, and conclusions or recommendations expressed in this material are those of the authors and do not necessarily reflect the views of the NSF or the NIH.

## AUTHOR CONTRIBUTIONS

J.A.K. and E.J.K.: project conceptualization, experimental design, data acquisition and analysis, manuscript preparation; M.I.M., R.M.K., Q.X.L.: data acquisition. All authors have given approval to the final version of the manuscript.

