## Supplementary Information for "An Activity-Based Nanosensor for Minimally-Invasive Measurement of Protease Activity in Traumatic Brain Injury"

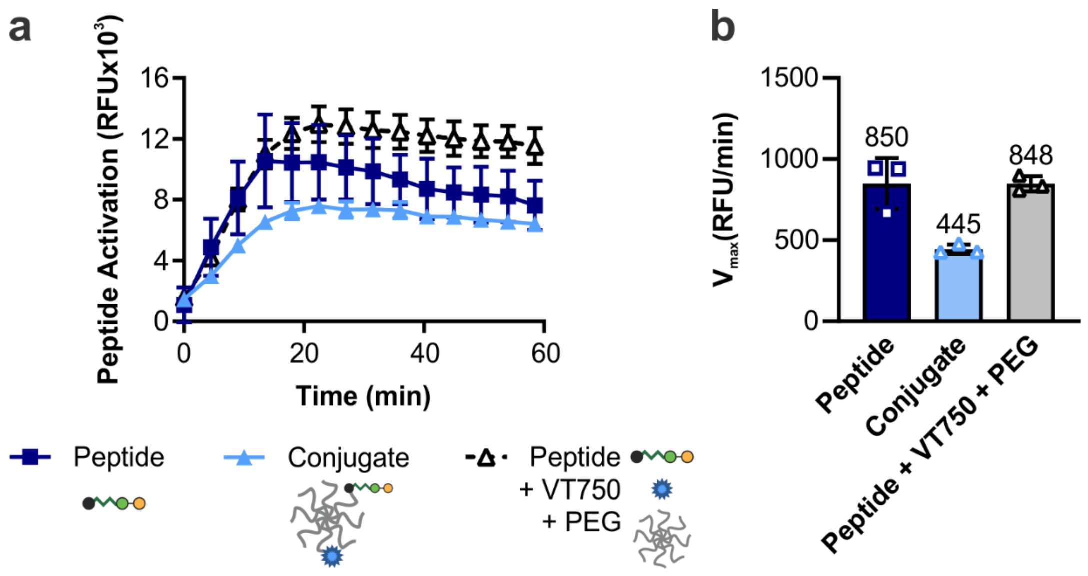


**Figure S1. Changes to peptide cleavage kinetics are due to conjugation of calpain substrate peptide to PEG. a.** Kinetic curves and **b.** $V_{max}$ at 8 µM peptide for calpain substrate peptide, TBI-ABN conjugate, and TBI-ABN conjugate components (n = 3 technical replicates, mean ± SD).

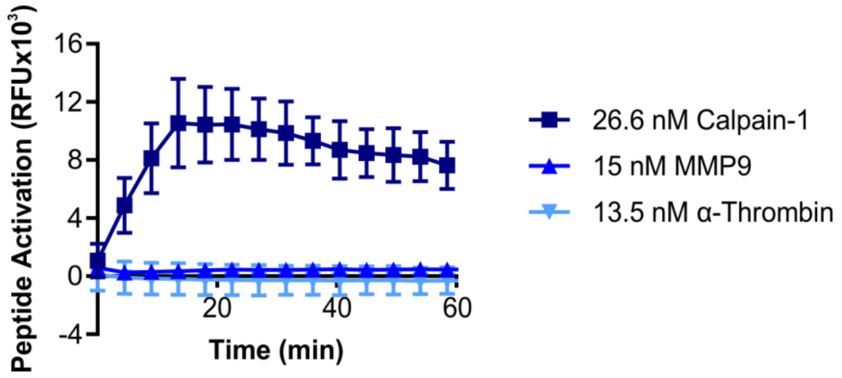


**Figure S2. Calpain substrate peptide is specifically cleaved by calpain-1.** Specificity assay of 8 µM calpain substrate peptide after incubation with calpain-1, MMP9, or α-thrombin (n = 3 technical replicates, mean ± SD).

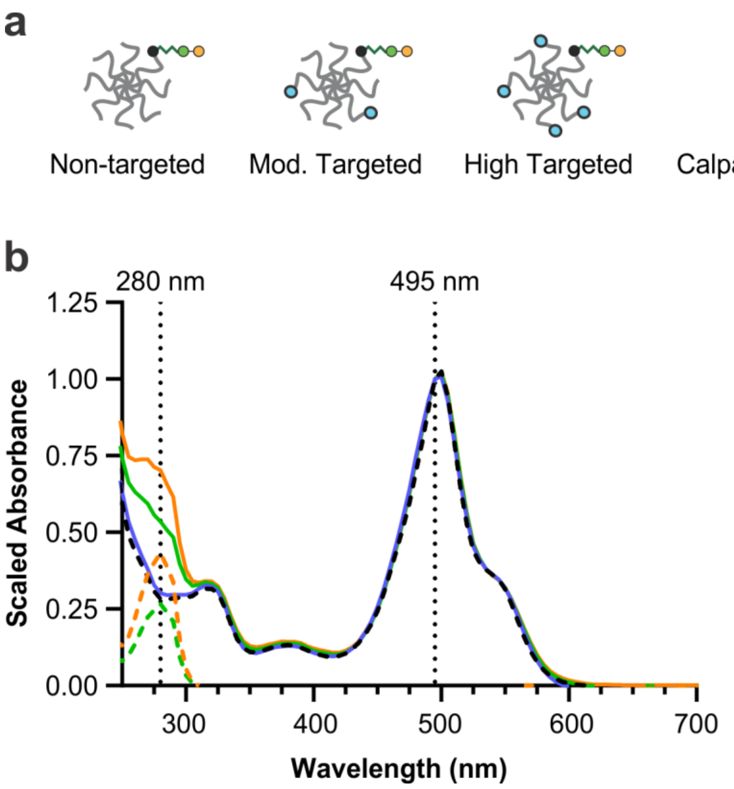


**Figure S3. Absorbance spectra of targeted TBI-ABNs in PBS show an increased absorbance at λ = 280 nm with the addition of HApep. a.** Schematic of targeted TBI-ABNs and peptide components. **b.** Scaled absorbance spectra of targeted TBI-ABNs and peptide components. Calpain substrate absorbance was measured by FAM absorbance at λ = 495 nm, and concentration was calculated from a standard of free peptide in PBS. HApep absorbance was measured by tryptophan content at λ = 280 nm, concentration was calculated with $\varepsilon_{280\ nm}$ = 5,500 $M^{-1}$ $cm^{-1}$ after correcting for the contribution to absorbance from calpain substrate.

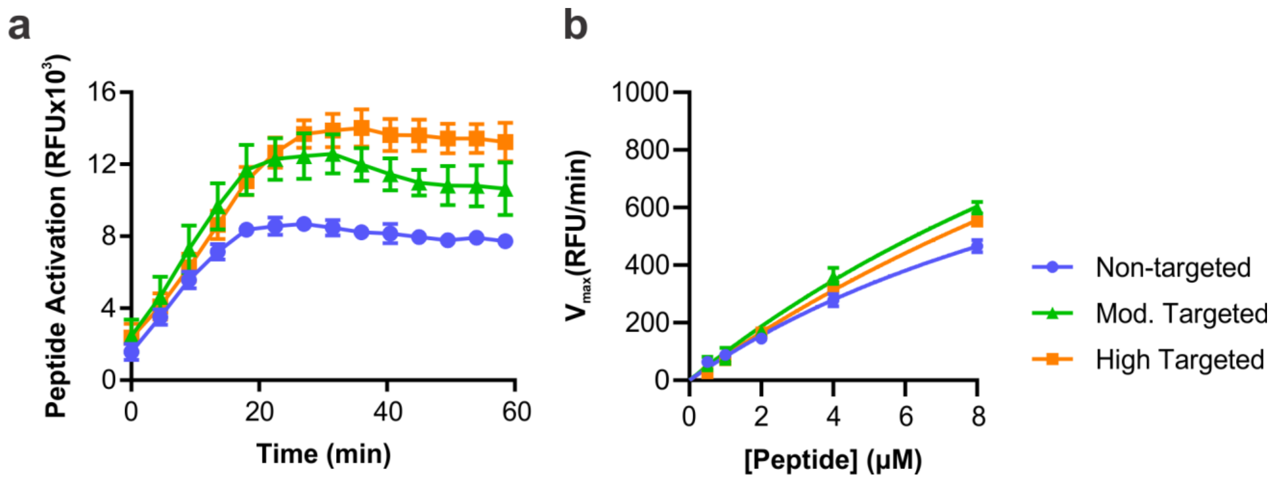


**Figure S4. Conjugation of HApep to PEG minimally changes the cleavage kinetics of calpain substrate peptide. a.** Kinetic curves at 8 µM calpain substrate peptide of targeted TBI-ABNs incubated with human calpain-1 (n = 3 technical replicates, mean ± SD). **b.** Michaelis Menten curves derived from the maximal velocities of cleavage (n = 3 technical replicates, mean ± SD).

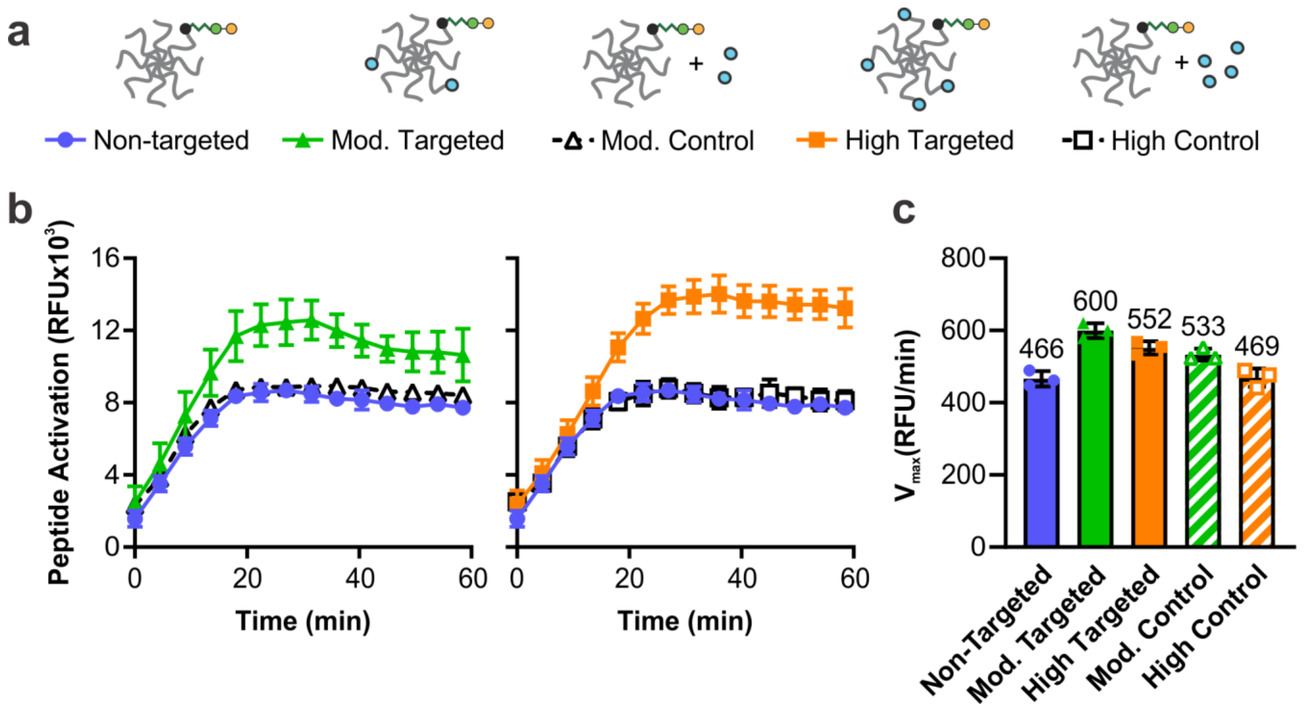


**Figure S5. Conjugation of HApep to PEG minimally changes the cleavage kinetics of calpain substrate peptide. a.** Non-targeted TBI-ABN was incubated with unconjugated HApep at equal ratios to the HApep in moderate and high targeted TBI-ABNs as controls for conjugation. **b.** Deconstructed conjugate control curves and **c.** bar graph of $V_{max}$ for moderate and high targeted conjugates at 8 µM calpain substrate peptide with human calpain-1 (n = 3 technical replicates, mean ± SD).

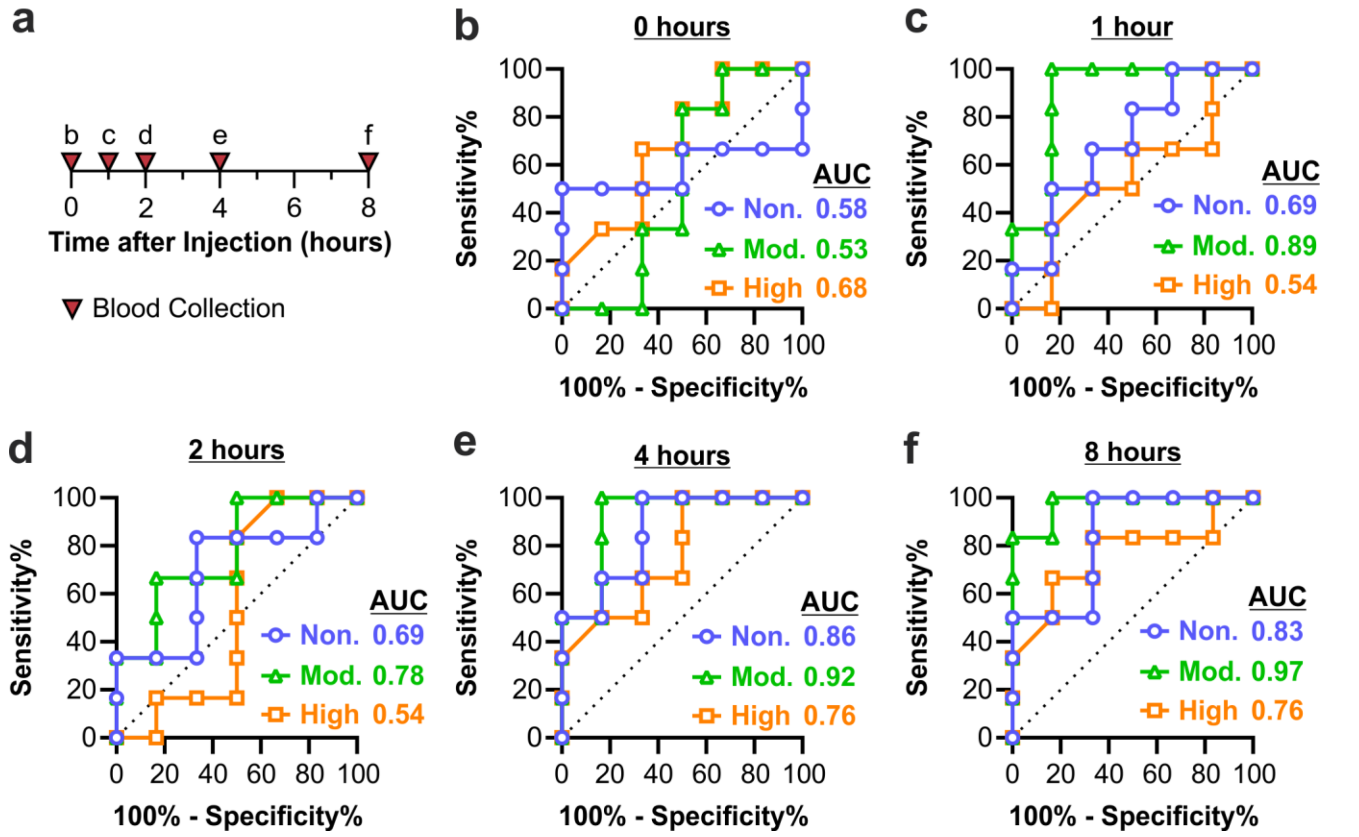


**Figure S6. Discrimination of injury via blood signal improves over time after injection in all targeting valencies. a.** Time points of blood collection and **b-f.** ROC curves classifying injured from uninjured mice using TBI-ABN fluorescence from blood samples at 0, 1, 2, 4, and 8 hours post-injection, respectively (n = 6). See **Table S1** for additional ROC statistics.

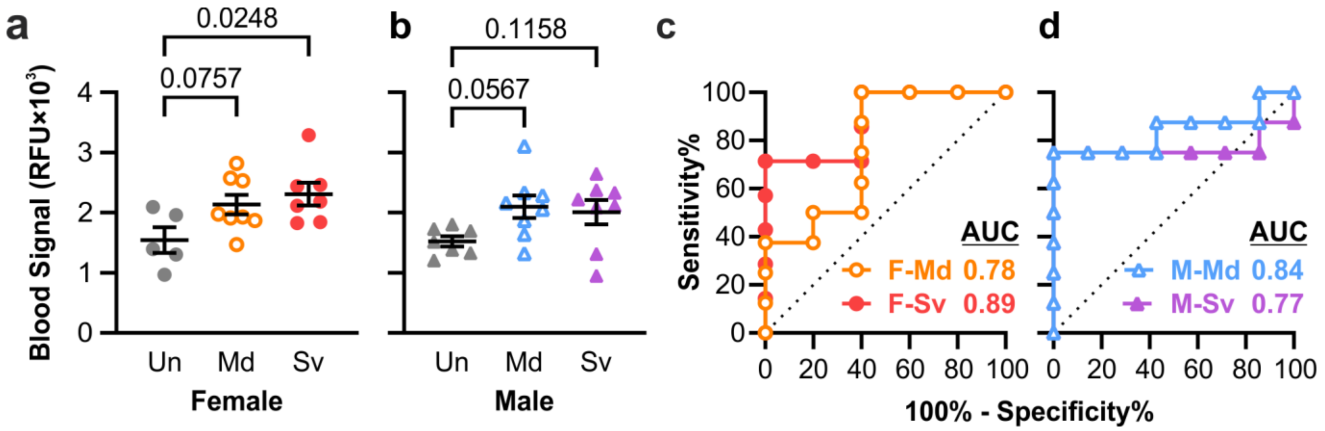


**Figure S7. TBI-ABN showed a slight increase in blood signal with injury across both sexes of mice. a, b.** TBI-ABN activation in blood at 4 hours post-injection as measured by fluorescence and **c, d.** corresponding ROC curves in female and male mice, respectively. (Un = uninjured; Md = mild CCI; Sv = severe CCI; F = female; M = male; n = 5 for F-Un; n = 7 for F-Sv and M-Un; n = 8 for F-Md, M-Md, and M-Sv; mean ± SE, ordinary one-way ANOVA with Dunnett's multiple comparisons post-hoc test against uninjured control). See **Table S2** for additional ROC statistics.

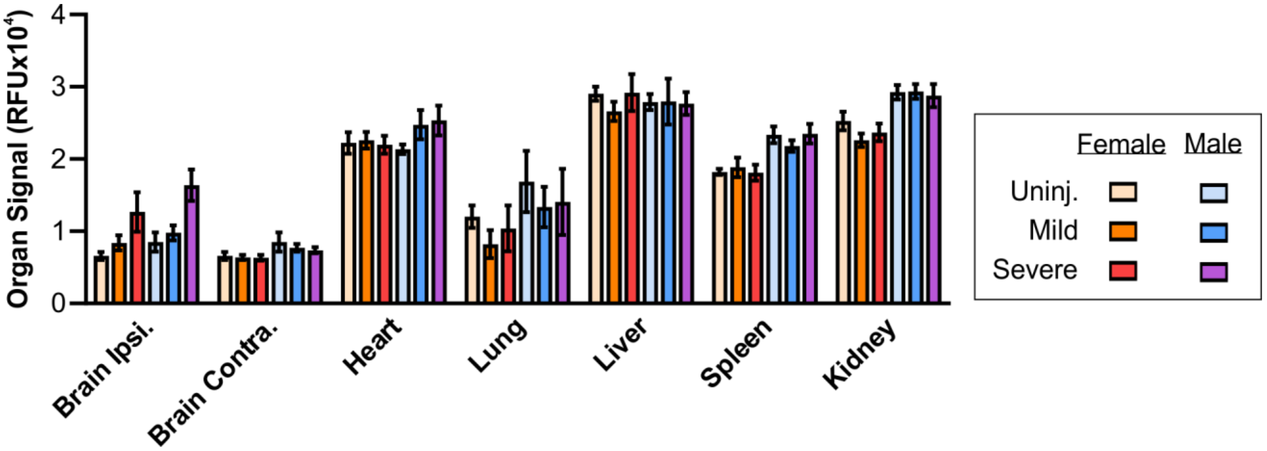


**Figure S8. TBI-ABN activation was comparable within injury groups across each organ and sex.** TBI-ABN activation in the brain (Ipsi. = injured cortical tissue, Contra. = contralateral cortical tissue, with Ipsi.+Contra. pooled for uninjured control) and in off-target organs in female and male mice with no, mild, or severe CCI (n = 5 for female uninjured; n = 7 for female severe and male uninjured; n = 8 for female mild, male mild, and male severe; mean ± SE).

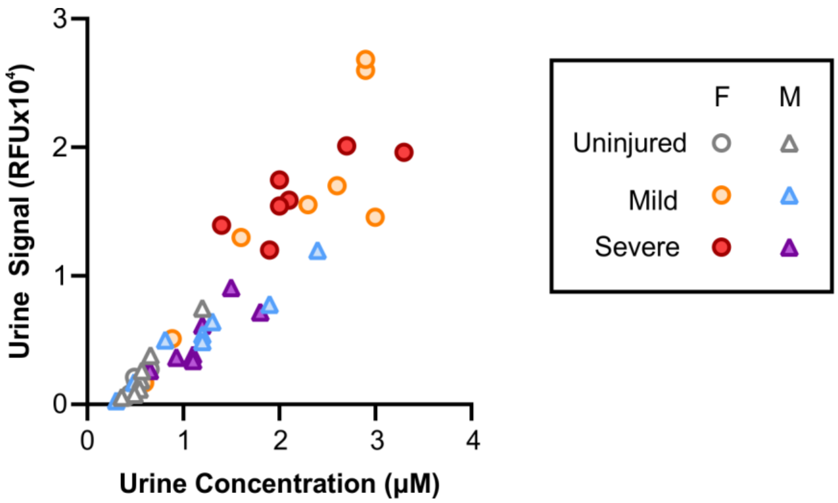


**Figure S9. ELISA measurements of c-Peptide concentration in urine samples correlate with fluorescence measurements of c-Peptide activation.** Scatterplot, fluorescence vs. ELISA concentration of cleaved TBI-ABN in urine collected at 1 hour post-injection (n = 43, Pearson correlation analysis: r = 0.9313, p<0.0001).

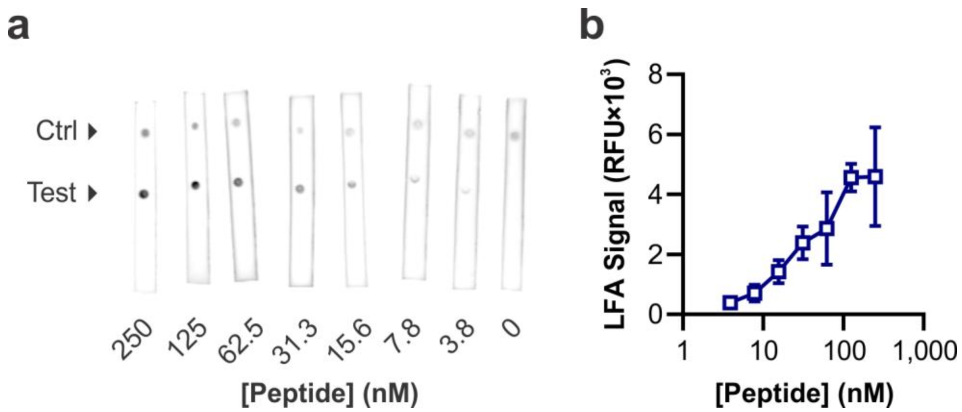


**Figure S10. Calpain substrate peptide can be quantified via LFA. a.** Representative fluorescent scans of LFAs for a standard of free calpain substrate peptide spiked into diluted mouse urine (n = 3 technical replicates, ctrl = control). **b.** Integrated fluorescence signal from the LFA test spots shows an increase in test spot signal with increasing peptide concentration (n = 3 technical replicates, mean ± SD).

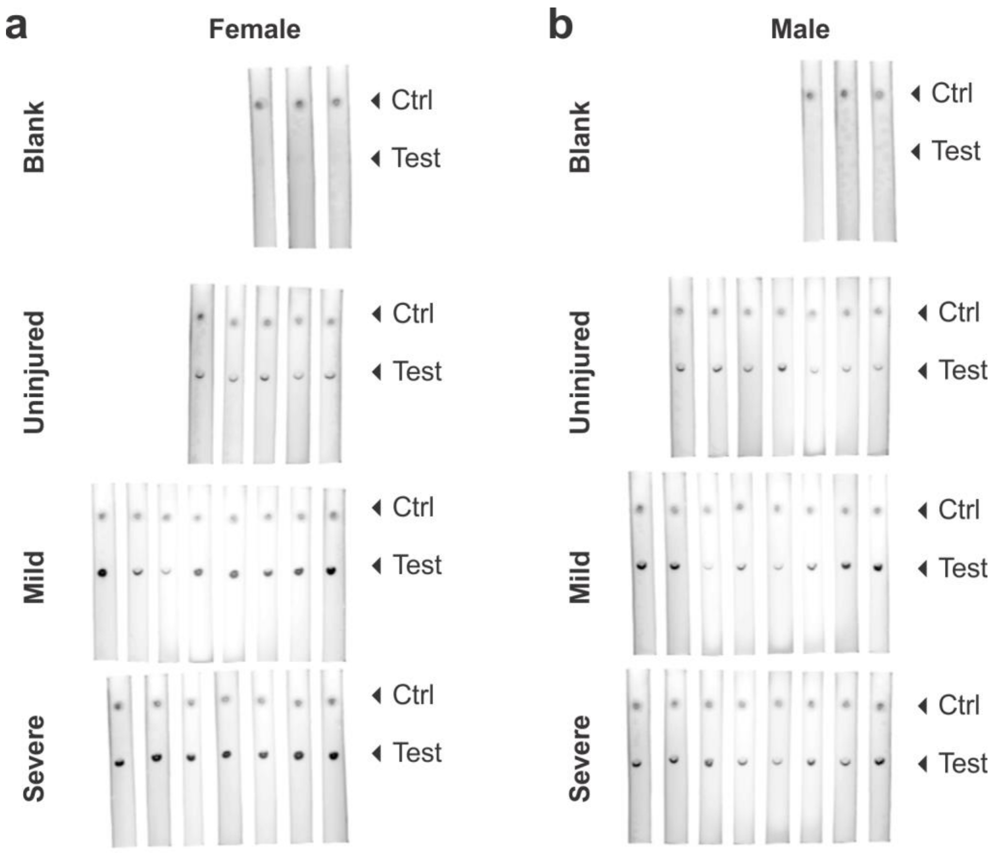


**Figure S11**. **LFAs can detect c-Peptide from in mouse urine samples. a, b.** Fluorescence scans of LFAs for urine samples taken from female and male mice in **Figure 5a**, respectively. All LFAs were imaged concurrently with the same imaging settings, and are cropped for clarity (Blank = sex-matched urine samples from uninjured mice that did not receive TBI-ABN, ctrl = control).

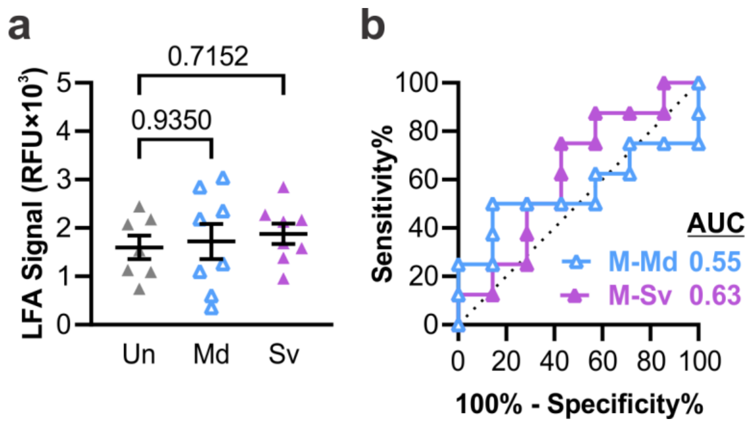


**Figure S12. Measurement of c-Peptide in the urine of male mice via LFA. a.** Accumulation of c-Peptide in urine at 1 hour post-injection as measured by integrated test spot signal from each LFA and **b.** corresponding ROC curves in male mice (Un = uninjured; Md = mild CCI; Sv = severe CCI; M = male; n = 7 for M-Un; n = 8 for M-Md and M-Sv; mean ± SE, ordinary one-way ANOVA with Dunnett's multiple comparisons post-hoc test against uninjured control). See **Table S2** for additional ROC statistics.

**Table S1. ROC Statistics, TBI-ABN Targeting Comparison**

| Figure | Measurement | TBI-ABN | Time Post-Injection | AUC | 95% CI | SE | P Value |
|---|---|---|---|---|---|---|---|
| 4c, S6 | TBI-ABN, Blood, Fluorescence | Non-Targeted | 0 hr | 0.58 | 0.21-0.95 | 0.19 | 0.6310 |
| | | | 1 hr | 0.69 | 0.38-1.00 | 0.16 | 0.2623 |
| | | | 2 hr | 0.69 | 0.38-1.00 | 0.16 | 0.2623 |
| | | | 4 hr | 0.86 | 0.65-1.00 | 0.11 | 0.0374 |
| | | | 8 hr | 0.83 | 0.59-1.00 | 0.12 | 0.0547 |
| | | Mod. Targeted | 0 hr | 0.53 | 0.16-0.89 | 0.19 | 0.8728 |
| | | | 1 hr | 0.89 | 0.67-1.00 | 0.11 | 0.0250 |
| | | | 2 hr | 0.78 | 0.50-1.00 | 0.14 | 0.1093 |
| | | | 4 hr | 0.92 | 0.74-1.00 | 0.089 | 0.0163 |
| | | | 8 hr | 0.97 | 0.89-1.00 | 0.043 | 0.0065 |
| | | High Targeted | 0 hr | 0.68 | 0.37-0.99 | 0.16 | 0.2980 |
| | | | 1 hr | 0.54 | 0.20-0.89 | 0.18 | 0.8102 |
| | | | 2 hr | 0.54 | 0.17-0.91 | 0.19 | 0.8102 |
| | | | 4 hr | 0.76 | 0.49-1.00 | 0.14 | 0.1282 |
| | | | 8 hr | 0.76 | 0.48-1.00 | 0.15 | 0.1282 |
| 4e | TBI-ABN, Urine, Fluorescence | Non-Targeted | 1 hr | 0.83 | 0.59-1.00 | 0.12 | 0.0547 |
| | | Mod. Targeted | 1 hr | 1.00 | 1.00-1.00 | 0.00 | 0.0039 |
| | | High Targeted | 1 hr | 0.76 | 0.47-1.00 | 0.15 | 0.1282 |

**Table S2. ROC Statistics, Sex- and Severity-Based Diagnostic Comparison**

| Figure | Measurement | Sex | Severity | AUC | 95% CI | SE | P Value |
|---|---|---|---|---|---|---|---|
| 5d | TBI-ABN, Urine, Fluorescence | Female | Mild | 0.93 | 0.77-1.00 | 0.08 | 0.0128 |
| | | | Severe | 1.00 | 1.00-1.00 | 0.00 | 0.0045 |
| 5e | TBI-ABN, Urine, Fluorescence | Male | Mild | 0.73 | 0.46-1.00 | 0.14 | 0.1325 |
| | | | Severe | 0.80 | 0.55-1.00 | 0.13 | 0.0491 |
| 5h | GFAP, Blood, ELISA | Female | Mild | 0.71 | 0.41-1.00 | 0.16 | 0.2232 |
| | | | Severe | 1.00 | 1.00-1.00 | 0.00 | 0.0045 |
| 5i | GFAP, Blood, ELISA | Male | Mild | 0.67 | 0.37-0.96 | 0.15 | 0.3017 |
| | | | Severe | 1.00 | 1.00-1.00 | 0.00 | 0.0019 |
| 6c | TBI-ABN, Urine, ELISA | Female | Mild | 0.98 | 0.90-1.00 | 0.039 | 0.0054 |
| | | | Severe | 1.00 | 1.00-1.00 | 0.00 | 0.0045 |
| 6d | TBI-ABN, Urine, ELISA | Male | Mild | 0.73 | 0.45-1.00 | 0.14 | 0.1325 |
| | | | Severe | 0.87 | 0.66-1.00 | 0.10 | 0.0177 |
| 6h | TBI-ABN, Urine, LFA | Female | Mild | 0.68 | 0.37-0.98 | 0.15 | 0.3055 |
| | | | Severe | 1.00 | 1.00-1.00 | 0.00 | 0.0045 |
| S7c | TBI-ABN, Blood, Fluorescence | Female | Mild | 0.78 | 0.49-1.00 | 0.15 | 0.1073 |
| | | | Severe | 0.89 | 0.69-1.00 | 0.10 | 0.0284 |
| S7d | TBI-ABN, Blood, Fluorescence | Male | Mild | 0.84 | 0.62-1.00 | 0.11 | 0.0279 |
| | | | Severe | 0.77 | 0.48-1.00 | 0.14 | 0.0826 |
| S12b | TBI-ABN, Urine, LFA | Male | Mild | 0.55 | 0.24-0.86 | 0.16 | 0.7285 |
| | | | Severe | 0.63 | 0.33-0.92 | 0.15 | 0.4179 |

**Table S3. Summary of Statistical Tests for In Vivo Experiments**

| Figure | Test | n (biological replicates) | F or t Value | P Value |
|---|---|---|---|---|
| 4b, non. | Two-way RM ANOVA with Geisser-Greenhouse correction and Sidak's multiple comparisons post-hoc test for each time point | n = 6 | F = 3.879 (4, 40), time x injury | p = n.s., all times |
| 4b, mod. | Two-way RM ANOVA with Geisser-Greenhouse correction and Sidak's multiple comparisons post-hoc test for each time point | n = 6 | F = 2.113 (4, 40), time x injury | p = n.s., 0, 1, and 2 hrs<br>p = 0.0283, 4 hr<br>p = 0.0477, 8 hr |
| 4b, high | Two-way RM ANOVA with Geisser-Greenhouse correction and Sidak's multiple comparisons post-hoc test for each time point | n = 6 | F = 2.022 (4, 40), time x injury | p = n.s., all times |
| 4d, non. | Unpaired t-test, two-tailed | n = 6 | t = 2.478, df = 10 | p = 0.0326 |
| 4d, mod. | Unpaired t-test, two-tailed | n = 6 | t = 6.256, df = 10 | p < 0.0001 |
| 4d, high | Unpaired t-test, two-tailed | n = 6 | t = 1.522, df = 10 | p = 0.1590 |
| 5b | Ordinary one-way ANOVA with Dunnett's multiple comparisons post-hoc test against uninjured control | n = 5, uninj.<br>n = 7, severe<br>n = 8, mild | F = 10.54 (2, 17) | p = 0.0019, mild<br>p = 0.0011, severe |
| 5c | Ordinary one-way ANOVA with Dunnett's multiple comparisons post-hoc test against uninjured control | n = 7, uninj.<br>n = 8, mild and severe | F = 1.966 (2, 20) | p = 0.1251, mild<br>p = 0.2526, severe |
| 5f | Ordinary one-way ANOVA with Dunnett's multiple comparisons post-hoc test against uninjured control | n = 5, uninj.<br>n = 7, mild and severe | F = 10.38 (2,16) | p = 0.7622, mild<br>p = 0.0018, severe |
| 5g | Ordinary one-way ANOVA with Dunnett's multiple comparisons post-hoc test against uninjured control | n = 6, uninj.<br>n = 8, mild and severe | F = 30.84 (2,19) | p = 0.7482, mild<br>p < 0.0001, severe |
| 5j | Two-way ANOVA with Dunnett's multiple comparisons test compared to uninjured within each organ | n = 5, uninj.<br>n = 7, severe<br>n = 8, mild | F = 2.232 (12, 129), injury severity x organs | p < 0.0001, severe within ipsi. brain<br>p = n.s., all other comparisons |
| 5k | Two-way ANOVA with Dunnett's multiple comparisons test compared to uninjured within each organ | n = 7, uninj.<br>n = 8, mild and severe | F = 2.282 (12, 154), injury severity x organs | p < 0.0001, severe within ipsi. brain<br>p = n.s., all other comparisons |
| 6a | Ordinary one-way ANOVA with Dunnett's multiple comparisons post-hoc test against uninjured control | n = 5, uninj.<br>n = 7, severe<br>n = 8, mild | F = 9.985 (2, 17) | p = 0.0020, mild<br>p = 0.0015, severe |
| 6b | Ordinary one-way ANOVA with Dunnett's multiple comparisons post-hoc test against uninjured control | n = 7, uninj.<br>n = 8, mild and severe | F = 2.834 (2, 20) | p = 0.0664, mild<br>p = 0.1275, severe |

| | | | | |
|---|---|---|---|---|
| 6g | Ordinary one-way ANOVA with Dunnett's multiple comparisons post-hoc test against uninjured control | n = 5, uninj.<br>n = 7, severe<br>n = 8, mild | F = 4.268 (2,17) | p = 0.3693, mild<br>p = 0.0195, severe |
| S7a | Ordinary one-way ANOVA with Dunnett's multiple comparisons post-hoc test against uninjured control | n = 5, uninj.<br>n = 7, severe<br>n = 8, mild | F = 3.987 (2, 17) | p = 0.0757, mild<br>p = 0.0248, severe |
| S7b | Ordinary one-way ANOVA with Dunnett's multiple comparisons post-hoc test against uninjured control | n = 7, uninj.<br>n = 8, mild and severe | F = 3.024 (2, 20) | p = 0.0567, mild<br>p = 0.1158, severe |
| S12a | Ordinary one-way ANOVA with Dunnett's multiple comparisons post-hoc test against uninjured control | n = 7, uninj.<br>n = 8, mild and severe | F =0.2389 (2, 20) | p = 0.9350, mild<br>p = 0.7152, severe |
